# Pregnancy associated plasma protein-aa regulates endoplasmic reticulum-mitochondria associations

**DOI:** 10.1101/2020.06.09.142505

**Authors:** Mroj Alassaf, Mary Halloran

**Author notes:** Corresponding author: Mary Halloran, Department of Integrative Biology, 1117 West Johnson Street, Madison, WI 53706.

## Abstract

The endoplasmic reticulum (ER) and mitochondria form close physical associations to facilitate calcium transfer, thereby regulating mitochondrial function and dynamics. For neurons with high metabolic demands, such as sensory hair cells, precise regulation of ER-mitochondria associations is especially critical for cell survival. We previously identified the secreted metalloprotease Pregnancy associated plasma protein-aa (Pappaa) as a novel regulator of mitochondrial function in zebrafish lateral line hair cells (Alassaf et al., 2019). Here, we show that *pappaa* mutant hair cells exhibit excessive and abnormally close ER-mitochondria associations, suggesting increased ER-mitochondria calcium transfer. Indeed, we find that *pappaa* mutant hair cells are more vulnerable to pharmacological induction of ER-calcium release. Additionally, *pappaa* mutant hair cells display ER stress and dysfunctional downstream processes of the ER-mitochondria axis including mitochondrial fragmentation and autophagy. Together our results support a model in which Pappaa regulates mitochondrial function, at least in part, by regulating ER-mitochondria associations.

## Introduction

Neuronal survival is critically dependent on proper mitochondrial function, which is regulated in part by close associations between mitochondria and the endoplasmic reticulum (ER). Despite the diverse pathologies of neurodegenerative diseases, increasing evidence suggests that a disrupted ER-mitochondria connection is a common underlying feature (Area-Gomez et al., 2012; Calì et al., 2013; Hedskog et al., 2013; Bernard-Marissal et al., 2015). Indeed, many of the cellular processes associated with neurodegeneration such as mitochondrial dysfunction, mitochondrial fragmentation, disrupted autophagy, and oxidative stress are regulated by the ER-mitochondria axis (Friedman et al., 2011; Rowland and Voeltz, 2012; Csordás et al., 2018).

Precise regulation of mitochondrial calcium levels is essential for mitochondrial metabolism and cell survival (Cárdenas et al., 2010). The association of mitochondria with ER facilitates efficient uptake of calcium by the mitochondria (Rizzuto et al., 1998; Krols et al., 2016). Calcium released from the ER forms highly concentrated calcium microdomains that are necessary to override the low affinity of the mitochondrial calcium uniporter (MCU), the primary mitochondrial calcium uptake channel (Rizzuto et al., 2012). However, overly close inter-organelle distances or excessive ER-mitochondria contact points can result in increased calcium transfer, which in turn can overstimulate mitochondrial bioenergetics leading to the production of pathological levels of the energy byproduct, reactive oxygen species (ROS). Accumulation of ROS can then lead to oxidative stress, fragmentation of the mitochondrial network, and cell death (Brookes et al., 2004; Deniaud et al., 2008; Adam-Vizi and Starkov, 2010; Esterberg et al., 2014; Iqbal and Hood, 2014; Ježek et al., 2018). Thus, tight regulation of the distance and frequency of contacts between the ER and mitochondria is critical for proper mitochondrial calcium load. Previous studies investigating regulation of ER-mitochondria associations have focused on identifying proteins involved in tethering the ER to the mitochondria. However, the upstream molecular pathways that act to prevent excessive and pathologically tight ER-mitochondria associations, and the mechanisms by which these molecular pathways are regulated remain unknown.

To investigate these mechanisms, we are using zebrafish lateral line hair cells, which share molecular, functional, and morphological similarities with mammalian inner ear hair cells. Hair cells of the inner ear and lateral line system are specialized sensory neurons that transduce acoustic information and relay it to the central nervous system (McPherson, 2018). Sensory hair cells are an excellent model to investigate mechanisms regulating ER-mitochondria associations. Their high metabolic requirements make them particularly vulnerable to disruptions in mitochondrial calcium load (Gonzalez-Gonzalez, 2017). Indeed, excess mobilization of calcium from the ER to the mitochondria was recently shown to underlie aminoglycoside-induced hair cell death in the zebrafish lateral line (Esterberg et al., 2014). These findings highlight the importance of the ER-mitochondria connection in regulating hair cell survival. Given that hair cell death is considered the primary cause of hearing loss (Eggermont, 2017), identifying molecular factors that regulate the ER-mitochondria connection may provide insight into potential therapeutic targets to combat hearing loss.

We showed previously that Pregnancy associated plasma protein-aa (Pappaa) regulates mitochondrial function in zebrafish lateral line hair cells. Pappaa is a locally secreted metalloprotease that regulates Insulin-like growth factor 1 (IGF1) signaling. IGF1 is sequestered extracellularly by IGF binding proteins (IGFBPs) that prevent it from binding to its receptor. Pappaa cleaves inhibitory IGFBs, thereby stimulating local IGF1 availability and signaling (Hwa et al., 1999). We found that loss of Pappaa, and the consequential reduction in IGF1 signaling, resulted in mitochondrial calcium overload, ROS buildup, and increased susceptibility to ROS-induced cell death (Alassaf et al., 2019). Moreover, other studies have shown that declining IGF1 levels correlate with conditions such as aging, neurodegenerative disorders, and obesity, in which the ER-mitochondria connection is known to be altered (Hedskog et al., 2013; Arruda et al., 2014; Liu and Zhu, 2017; Lee et al., 2018). However, a causative link between IGF1 signaling and ER-mitochondria associations has not been shown.

In this study, we reveal that Pappaa, an extracellular regulator of IGF1 signaling, is critical for regulation of the ER-mitochondria connection. We use electron microscopy (EM) to show that ER-mitochondria associations are closer and greater in number in *pappaa* mutant hair cells. Hair cells in *pappaa* mutants are more sensitive to pharmacological induction of ER-mediated calcium release and show increased mitochondrial fragmentation and stunted autophagic response. Loss of Pappaa also results in ER stress and activation of the unfolded protein response. Together, our results suggest that Pappaa exerts its effect on hair cell survival by serving as a key regulator of the ER-mitochondria axis. Given the multiple functions and ubiquitous expression of IGF1 receptors, a factor such as Pappaa that can locally activate IGF1 signaling may provide a more promising therapeutic target to prevent hearing loss.

## Results and Discussion

### Pappaa influences ER-mitochondria associations

Zebrafish lateral line hair cells lie on the surface of the skin and are arranged into structures called neuromasts (Figure 1A-B). Each neuromast consists of a cluster of hair cells surrounded by the glia-like support cells. Our previous work showed that hair cells in *pappaa* mutants (hereafter referred to as *pappaa*^*p170*^) are more susceptible to ROS-induced death and their mitochondria have increased calcium load (Alassaf et al., 2019). Therefore, we hypothesized that *pappaa*^*p170*^ mitochondria may have an increased frequency of ER-mitochondria associations. To test this, we used EM to visualize ER-mitochondria associations. We collected 80 nm-thick sections within head neuromasts and quantified the number of ER tubules within 100 nm of mitochondria, the maximum distance for effective mitochondrial calcium uptake (Csordás et al., 2018). We found that *pappaa*^*p170*^ hair cells had on average a greater number of ER tubules in close association with each mitochondrion (Figure 1C-D). Furthermore, we quantified the percentage of mitochondria in close association with a given number of ER tubules and found that more mitochondria in *pappaa*^*p170*^ hair cells were in close association with multiple ER tubules compared to wild type hair cells (Figure 1C, E). These results suggest that Pappaa regulates the frequency of ER-mitochondria associations.

**Figure 1.**
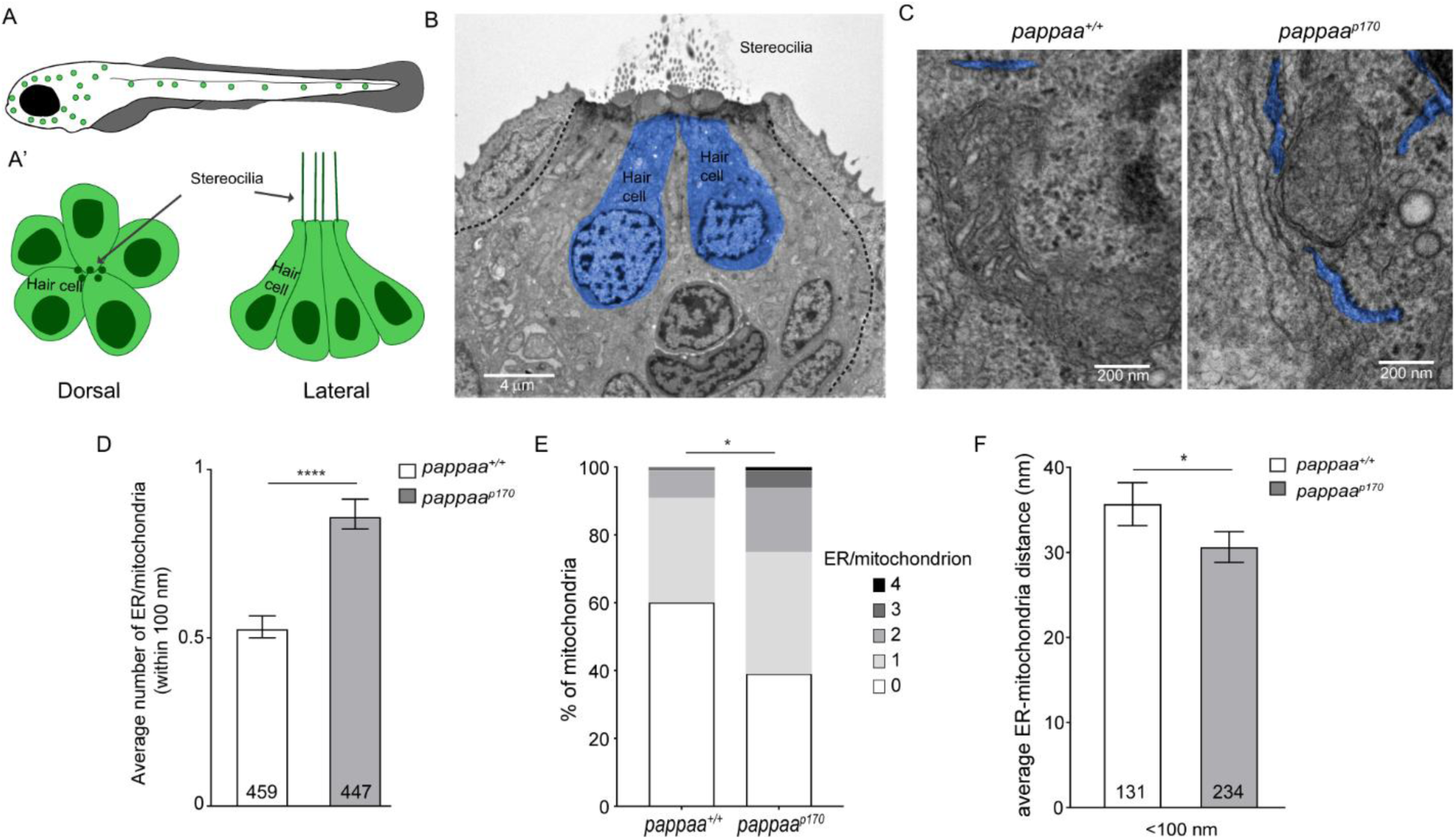
Pappaa regulates ER-mitochondria associations. (A-A’) Schematic of zebrafish lateral line hair cells. (B) Representative EM image of lateral line neuromast (dashed) showing lateral line hair cells (blue) in 5 dpf larva. (C) Representative EM images of ER-mitochondria associations in wild type and *pappaa* hair cells. (D) Mean number of ER tubules within 100 nm of mitochondria. \*\*\*\**p*<0.0001 *t* test, Mann-Whitney correction. N= 459 mitochondria (wild type) and 447 mitochondria (*pappaa*^*p170*^) collected from 6 larvae/genotype. Error bars=SEM. (E) Percentages of mitochondria associated with 0, 1, 2, 3, or 4 ER tubules. \**p*<0.05 Chi-square test. N= 459 mitochondria (wild type) and 447 mitochondria (*pappaa*^*p170*^) collected from 6 larvae/genotype. (F) Mean distance between the ER and mitochondria that are within 100 nm of each other. *\*p<*0.05 *t* test, Mann-Whitney correction. N= 131 ER-mitochondria associations (wild type) and 234 ER-mitochondria associations (*pappaa*^*p170*^) collected from 6 larvae/genotype. Error bars=SEM. **Figure 1-source data 1**. Mean number of ER tubules/mitochondrion **Figure 1-source data 2**. Percentages of mitochondria that are associated with 0, 1, 2, 3, or 4 ER tubules. **Figure 1-source data 3**. Mean ER-mitochondria distance.

The distance between ER and mitochondria also influences the efficiency of mitochondrial calcium intake and must be tightly regulated (Rizzuto et al., 2012). Thus, we asked whether *pappaa*^*p170*^ hair cells also had tighter ER-mitochondria associations. We measured the distances between all ER and mitochondria within 100 nm of each other. We found that *pappaa*^*p170*^ hair cells had shorter ER-mitochondria distance on average (Figure 1C, F). Taken together, our data suggest that Pappaa is important for preventing excessive and overly tight ER-mitochondria associations. Interestingly, another study used a synthetic linker to shorten the inter-organelle distance and showed a surge in mitochondria calcium load (Csordás et al., 2006), suggesting that the excessively tight connections we see in *pappaa*^*p170*^ hair cells could be causing their increased mitochondrial calcium load.

Although we previously reported no difference in mitochondrial mass between wild type and *pappaa*^*p170*^ hair cells (Alassaf et al., 2019), it remains possible that the increase in ER-mitochondria associations in *pappaa*^*p170*^ is a result of an overabundance of ER. Indeed, there is evidence to suggest that the ER undergoes expansion and heightened biogenesis under oxidative stress (Urra and Hetz, 2012). If that were the case in *pappaa*^*p170*^ hair cells, it could indicate that Pappaa functions to prevent oxidative stress by attenuating the buildup of mitochondrial ROS and consequently preventing ER expansion. Alternatively, the shortened distance between the ER and mitochondria may have initially triggered mitochondrial dysfunction leading to oxidative stress and ER expansion. Regardless, our results showing *pappaa*^*p170*^ hair cells also have a shortened inter-organelle distance suggest a role for Pappaa in actively regulating these interactions.

### Pappaa loss sensitizes hair cells to increased ER-mitochondria calcium transfer

Given the increased ER-mitochondria associations in *pappaa*^*p170*^ hair cells, we hypothesized that they experience augmented ER-mitochondria calcium transfer. If so, we would expect that *pappaa*^*p170*^ hair cells would show increased vulnerability to pharmacological stimulation of ER-mitochondria calcium transfer. We used two pharmacological approaches to increase calcium transfer. First, we treated 5 days post fertilization (dpf) wild type and *pappaa*^*p170*^ larvae with adenophostin A, an inositol 1,4,5-trisphosphate receptor (IP3R) agonist, to induce calcium efflux from the ER. Second, we treated larvae with Thapsigargin, a sarco/endoplasmic reticulum Ca^2+^-ATPase (SERCA) inhibitor, which prevents ER calcium influx and thereby allows for more available calcium in the cytosol to be taken up by the mitochondria (Figure 2A) (Esterberg et al., 2014). To assess hair cell survival, we used *Tg(brn3c:mGFP)* larvae, in which the lateral line hair cells are labeled with membrane-targeted GFP. We found that treatment for 1 hour with either drug at doses that are not cytotoxic to wild type hair cells resulted in a marked reduction in hair cell survival in *pappaa*^*p170*^ larvae (Figure 2B-C). These findings suggest that *pappaa*^*p170*^ hair cells are more sensitive to fluctuations in calcium and further support the idea that *pappaa*^*p170*^ hair cells have a disrupted ER-mitochondria connection that results in greater mitochondrial calcium uptake. However, it is difficult to discern whether the elevated levels of calcium in *pappaa*^*p170*^ mitochondria are solely due to the altered ER-mitochondria associations. It is possible that *pappaa*^*p170*^ hair cells experience increased ER calcium efflux and/or mitochondrial calcium uptake. Protein kinase B (Akt), which is a downstream effector of multiple signaling pathways including IGF1, has been shown to inhibit IP3Rs and reduce ER calcium release in HeLa cells (Marchi et al., 2008). If Akt has a similar role in hair cells, it is possible that Pappaa-IGF1 signaling may normally act to increase Akt-induced suppression of IP3Rs leading to reduced ER calcium release. Further investigation into the activity of ER and mitochondria associated calcium channels in *pappaa*^*p170*^ hair cells is needed to answer this question.

**Figure 2.**
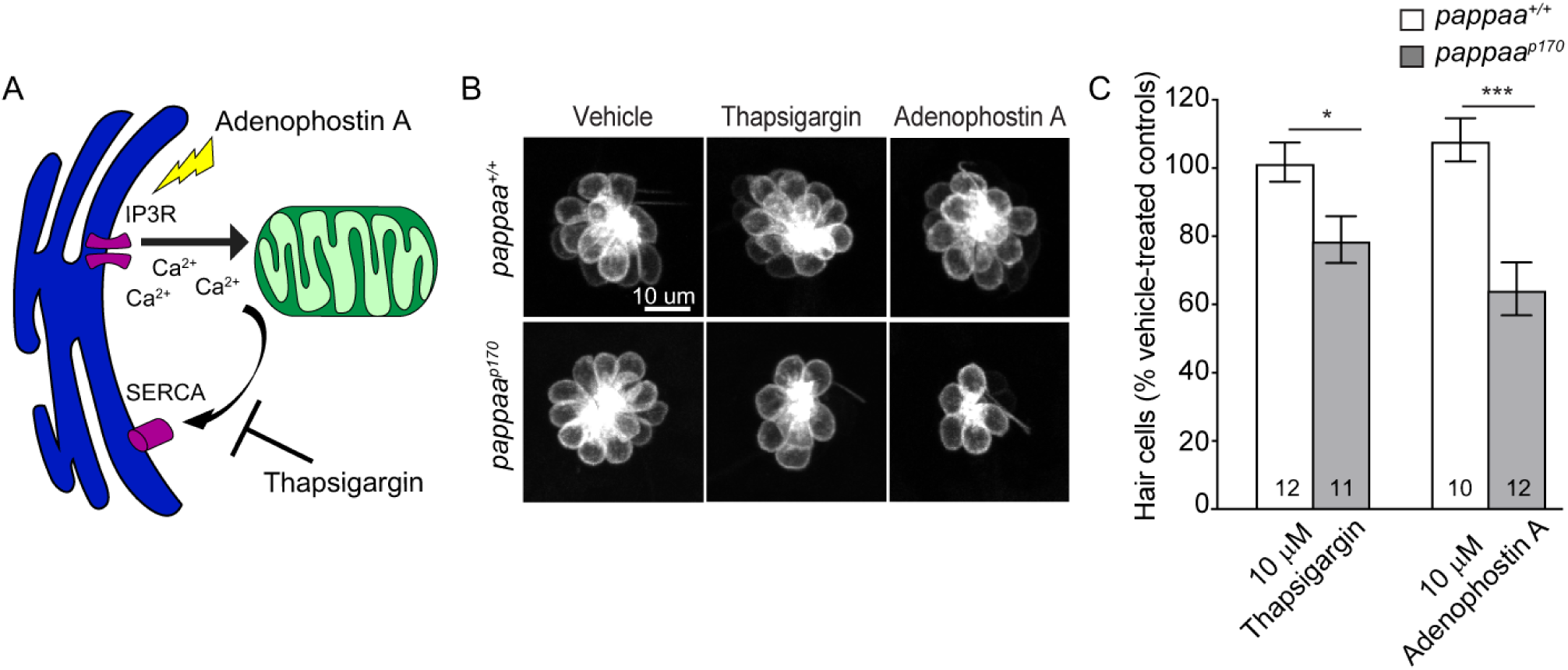
*pappaa*^*p170*^ hair cells are more sensitive to disruption in ER-mitochondria calcium signaling. (A) Schematic of calcium channel modulators. Thapsigargin inhibits calcium uptake by the ER by blocking the SERCA pump. Adenophostin A stimulates calcium release through activation of the IP3Rs. (B) Representative images of *brn3c:mGFP* labeled hair cells from vehicle, 10 µM Thapsigargin, or 10 µM Adenophostin A treated larvae. (C) Mean percentage of surviving hair cells following a 1-hour treatment with either Thapsigargin or Adenosphostin A at 5 dpf. To calculate hair cell survival percentage, hair cell number 4 hours post-drug treatment was normalized to mean hair cell number in vehicle treated larvae of the same genotype. \**p*<0.05, \*\*\**p*<0.001, two-way ANOVA, Holm-Sidak post test. N = 10-12 larvae per group (shown at base of bars), 3 neuromasts/ larva were analyzed. **Figure 2-source data 1**. Hair cell survival following treatment with Thapsigargin and Adenophostin A in wild type and *pappaa*^*p170*^ larvae.

### Pappaa loss causes mitochondrial fragmentation

Mitochondria undergo constant fission and fusion as a form of quality control. Damaged mitochondria are excised from the mitochondrial network through the process of fission and cleared by autophagy, while the remaining reusable pool of mitochondria fuse back together (Ni et al., 2015). Thus, a balance between mitochondrial fission and fusion must be achieved to maintain cellular homeostasis (Liesa et al., 2009). Under pathological conditions, such as oxidative stress in which there is an excess of damaged mitochondria, the balance between fission and fusion tips toward fission, and fragmentation of the mitochondrial network occurs ultimately leading to cell death. Recently, it was shown that mitochondrial fission occurs at the site of ER-mitochondria contacts. ER tubules were found to wrap around mitochondria and mark the site for fission (Friedman et al., 2011). Given that *pappaa*^*p170*^ hair cells experience oxidative stress, have dysfunctional mitochondria (Alassaf et al., 2019), and exhibit increased ER-mitochondria associations, we hypothesized that *pappaa*^*p170*^ hair cells experience increased fission. Mitochondrial morphology can be predictive of fission and fusion events. Small, round and clustered mitochondria are often indicative of fission events, whereas large and branched mitochondria are indicative of fusion events. An increase in fission is associated with fragmentation of the mitochondrial network. To test our hypothesis, we used EM and measured several parameters of mitochondrial morphology in wild type and *pappaa*^*p170*^ hair cells, including the area, perimeter, aspect ratio (major axis/minor axis; a measure of elongation), circularity (4π x Area/Perimeter^2^), and network interconnectivity (area/perimeter) (Wiemerslage and Lee, 2016). We found that *pappaa*^*p170*^ mitochondria had a smaller area, perimeter, and aspect ratio. Moreover, they were more circular and showed reduced network interconnectivity suggesting excessive mitochondrial fission and network fragmentation (Figure 3A-F) (Wiemerslage and Lee, 2016). To further investigate mitochondrial integrity in whole cells, we labeled hair cells with the vital mitochondrial dye Mitotracker. Using confocal microscopy, we acquired z-stacks of neuromast hair cells. The mitochondrial network in *pappaa*^*p170*^ hair cells appeared visibly fragmented compared to wild type (Figure 3G). Analysis of mitochondrial circularity using ImageJ (Figure 3H) revealed results similar to the EM data, supporting the idea that *pappaa*^*p170*^ mitochondria are fragmented. Taken together, our data suggest that Pappaa regulates mitochondrial fission, likely via ER-associated processes. Our findings are consistent with a previous study showing that treatment with IGF1 maintained mitochondrial integrity in denervated muscle cells by promoting mitochondrial fusion and preventing mitochondrial fission (Ding et al., 2017).

**Figure 3.**
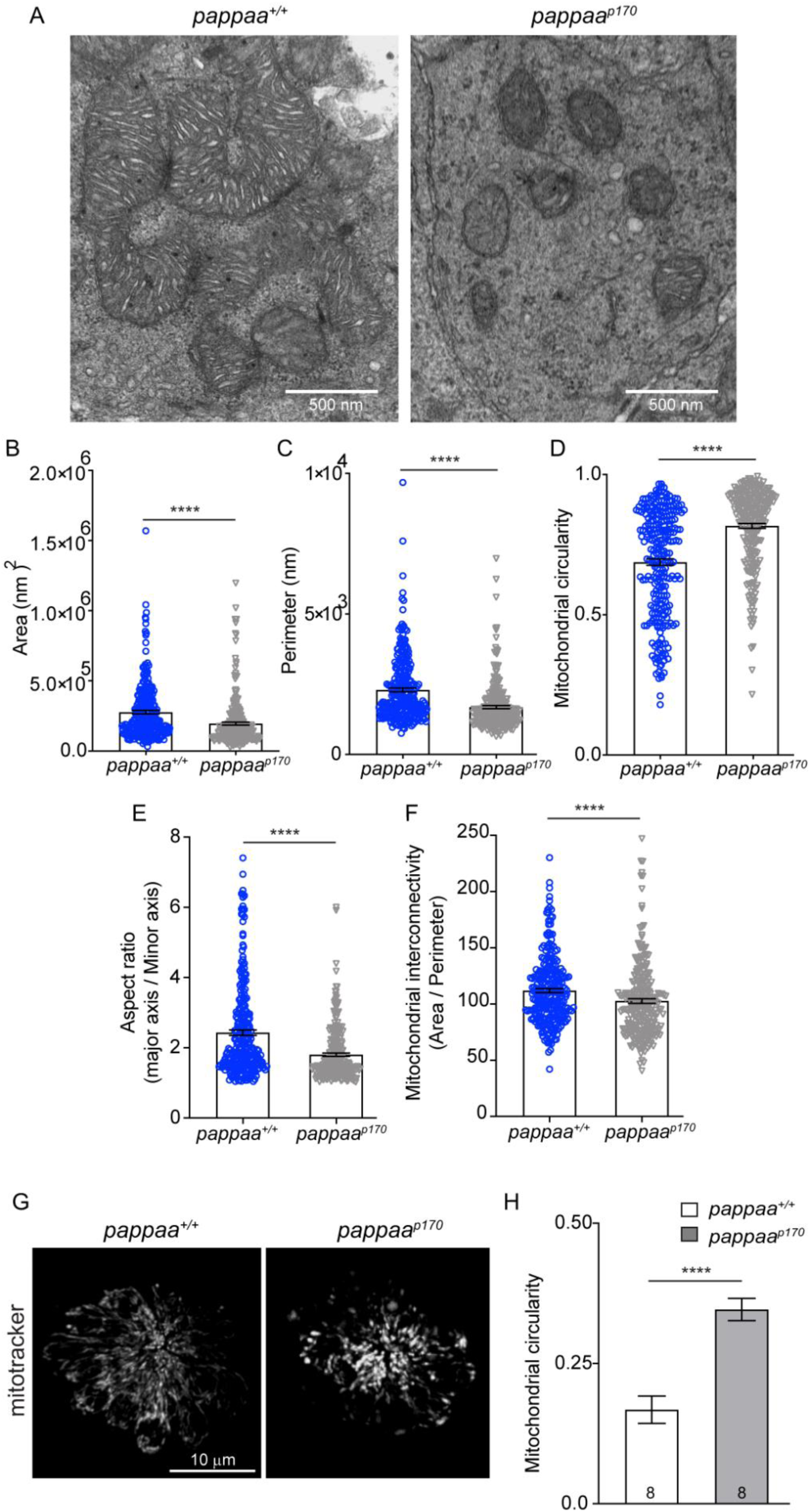
Pappaa loss causes mitochondrial fragmentation. (A) Representative EM images of mitochondria in lateral line hair cells in 5 dpf wild type and *pappaa*^*p170*^ larvae. (B-F) Mean mitochondrial (B) area, (C) perimeter, (D) circularity, (E) aspect ratio, and (F) interconnectivity in 5 dpf wild type and *pappaa*^*p170*^ lateral line hair cells. \*\*\*\**p*<0.0001 *t* test, Mann-Whitney correction. N= 272 mitochondria (wild type) and 262 mitochondria (*pappaa*^*p170*^) collected from 6 larvae/genotype. (G) Representative images of 5 dpf wild type and *pappaa*^*p170*^ lateral line hair cells loaded with the vital mitochondrial dye, Mitotracker. (H) Mean mitochondrial circularity measured from Z-stack max intensity projections of wild type and *pappaa*^*p170*^ lateral line hair cells. \*\*\*\**p<*0.0001 *t* test, Welch correction. N=8 larvae per group (shown at base of bars), 1 neuromast/ larva. Error bars=SEM. **Figure 3-source data 1**. Mitochondrial area in wild type and *pappaa*^*p170*^ lateral line hair cells. **Figure 3-source data 2**. Mitochondrial perimeter in wild type and *pappaa*^*p170*^ lateral line hair cells. **Figure 3-source data 3**. Mitochondrial circularity in wild type and *pappaa*^*p170*^ lateral line hair cells. **Figure 3-source data 4**. Mitochondrial aspect ratio in wild type and *pappaa*^*p170*^ lateral line hair cells. **Figure 3-source data 5**. Mitochondrial interconnectivity in wild type and *pappaa*^*p170*^ lateral line hair cells. **Figure 3-source data 6**. Mitochondrial circularity in wild type and *pappaa*^*p170*^ lateral line hair cells measured by mitotracker.

### Pappaa regulates neomycin-induced autophagy

We next asked if other processes known to be regulated by the ER-mitochondria axis are disrupted in *pappaa*^*p170*^ hair cells. Multiple studies suggest that autophagy, a bulk degradation process, is strongly influenced by the ER-mitochondria axis. Pharmacological and genetic manipulations that tighten or increase ER-mitochondria associations result in reduced autophagosome formation (Sano et al., 2012; Gomez-Suaga et al., 2017a; Gomez-Suaga et al., 2017b). Therefore, we asked whether *pappaa*^*p170*^ hair cells have an attenuated autophagic response. Interestingly, neomycin, to which *pappaa*^*p170*^ hair cells show hypersensitivity (Alassaf et al., 2019), was shown to be sequestered into autophagosomes upon entry into hair cells (Hailey et al., 2017; He et al., 2017). These authors also showed that blocking autophagy exacerbates neomycin-induced damage presumably due to the heightened exposure of vital intracellular components to neomycin. Thus, neomycin uptake into autophagosomes can be used as an indicator of the robustness of the autophagic response. To test whether *pappaa*^*p170*^ hair cells show a dampened autophagic response to neomycin treatment, we used a fluorescently tagged neomycin (Neomycin-Texas Red). Upon cell entry, neomycin-Texas Red appears as a diffused pool in the cytosol. Within minutes, neomycin gets captured by autophagosomes and high fluorescent intensity puncta begin to appear (Hailey et al., 2017) (Figure 4 and Video 1). We counted the number of puncta within visually accessible neuromast hair cells in wild type and *pappaa*^*p170*^ larvae. We found that *pappaa*^*p170*^ hair cells had fewer neomycin-Texas Red puncta (Figure 4 and Video 2), indicating that neomycin-induced autophagy was attenuated in *pappaa*^*p170*^ hair cells. This reduction in autophagy may allow neomycin to accumulate in the cytosol, causing more damage to intracellular compartments, and likely contributing to the increase in neomycin-induced hair cell death in *pappaa*^*p170*^ larvae (Alassaf et al., 2019).

**Figure 4.**
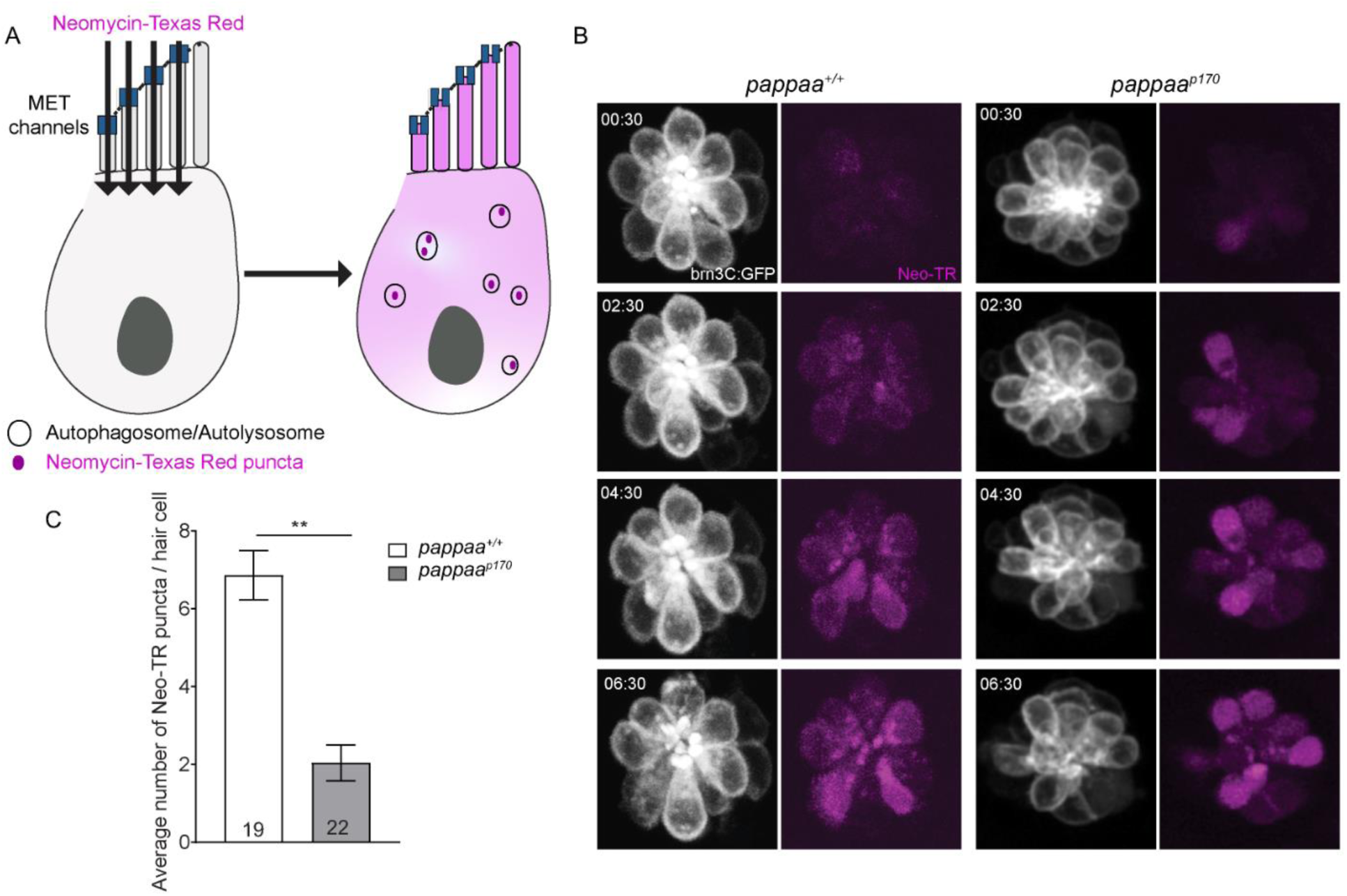
Pappaa regulates neomycin-induced autophagy. (A) Representative time lapse images of *brn3C:mGFP* labeled neuromast hair cells (white) at 5 dpf following exposure to 10 μM Neomycin-Texas Red (magenta). (B) mean number of Neomycin-Texas Red puncta/ hair cell in wild type and *pappaa*^*p170*^ larvae at 5 dpf. \*\**p*<0.01 *t* test, Mann-Whitney correction. N= 19 hair cells (wild type) and 22 hair cells (*pappaa*^*p170*^) collected from 4 larvae/genotype. Error bars=SEM. See videos 1 and 2. **Figure 4-source data 1**. Mean number of Neomycin-Texas Red puncta in wild type and *pappaa*^*p170*^ lateral line hair cells.

## Videos

**Video 1. Wild type lateral line hair cells following exposure to Neomycin-Texas red**. This video was constructed from time lapse images of wild type *brn3c:mGFP* lateral line hair cells (left) following exposure to 10 μM Neomycin-Texas red (right).

**Video 2. *pappaa***^***p170***^ **lateral line hair cells following exposure to Neomycin-Texas red**. This video was constructed from time lapse images of *pappaa*^*p170*^ *brn3c:mGFP* lateral line hair cells (left) following exposure to 10 μM Neomycin-Texas red (right).

### Pappaa loss causes ER stress

We next sought to define how Pappaa regulates ER-mitochondria associations. In addition to acting as the largest calcium reserve in the cell, the ER is the primary hub for protein processing. Newly synthesized proteins enter the ER for chaperone-assisted folding (Gregersen et al., 2006). Improper or insufficient folding results in the accumulation of unfolded proteins, which causes ER stress and increased calcium efflux (Houck et al., 2012). This efflux results in an increase in highly concentrated calcium “hot spots” that act as arresting signals for mitochondria, tethering the mitochondria to the ER and thereby increasing the frequency and tightness of ER-mitochondria associations (Bravo et al., 2011; Bravo et al., 2012). To ask whether *pappaa*^*p170*^ hair cells experience ER stress, we FACsorted hair cells from wild type and *pappaa*^*p170*^ *Tg(brn3c:mGFP)* larvae and evaluated gene expression of the unfolded protein response (UPR) by RT-qPCR. The UPR consists of three ER transmembrane receptors that act as sensors for unfolded proteins (Figure 5A). Normally, Bip, an ER-resident chaperone, is bound to UPR receptors. When unfolded proteins begin to accumulate, Bip dissociates from the UPR receptors to assist in protein folding. The uncoupling of Bip activates the UPR signaling cascade signifying ER stress. During the early phase of ER stress, the UPR promotes cell survival by improving the folding capacity of the ER through the upregulation of pro-survival factors including *bip, atf4*, and the splicing of *xbp1*. However, a switch from an adaptive to a pro-apoptotic UPR occurs during the late phase of ER stress, in which *chop*, a pro-apoptotic transcription factor, is upregulated. (Oslowski and Urano, 2011; Fu et al., 2015). Our analysis revealed that components of the “adaptive” UPR pathway (*bip, atf, and spliced-xbp1*) were upregulated in *pappaa*^*p170*^ hair cells (Figure 5B). Notably, the “pro-apoptotic” factor, c*hop*, was not differentially expressed in *pappaa*^*p170*^ hair cells. These results suggest that *pappaa*^*p170*^ hair cells experience early ER stress but not the later, pro-apoptotic response, which may explain the absence of spontaneous hair cell death in *pappaa*^*p170*^ larvae (Alassaf et al., 2019) To ask whether *pappaa*^*p170*^ hair cells are more vulnerable to pharmacological induction of protein unfolding given their already active UPR, we treated 5 dpf wild type and *pappaa*^*p170*^ *Tg(brn3c:mGFP)* larvae with tunicamycin, an inhibitor of glycoprotein biosynthesis (Oslowski and Urano, 2011), for 24 hours, and analyzed hair cell survival. *pappaa*^*p170*^ larvae showed a marked reduction in hair cell survival compared to wild type, suggesting that they are closer to the cytotoxic threshold of ER stress (Figure 5C). Together, these findings suggest that Pappaa may regulate ER-mitochondria associations by promoting ER homeostasis. It is important to note that the ER and mitochondria are engaged in a constant feedback loop. ER-mediated calcium release acts a key regulator of mitochondrial bioenergetics, and thereby, ROS production (Gorlach et al., 2015). Notably, this relationship is not unidirectional. Mitochondria-generated ROS can stimulate the activity of ER-associated calcium channels causing further calcium release (Brookes et al., 2004; Chaudhari et al., 2014; Gorlach et al., 2015). Given that many ER-resident molecular chaperones are calcium-dependent, ER stress can be both a cause and a consequence of mitochondrial dysfunction (Zhu and Lee, 2015). In future studies, it will be interesting to define Pappaa’s primary intracellular target, whether it was the mitochondria, ER, or both. Furthermore, given the novelty of Pappaa’s role in regulating ER-mitochondria associations, it will be interesting to identify the downstream targets of Pappaa-mediated IGF1 signaling within the ER-mitochondria tethering complex.

**Figure 5.**
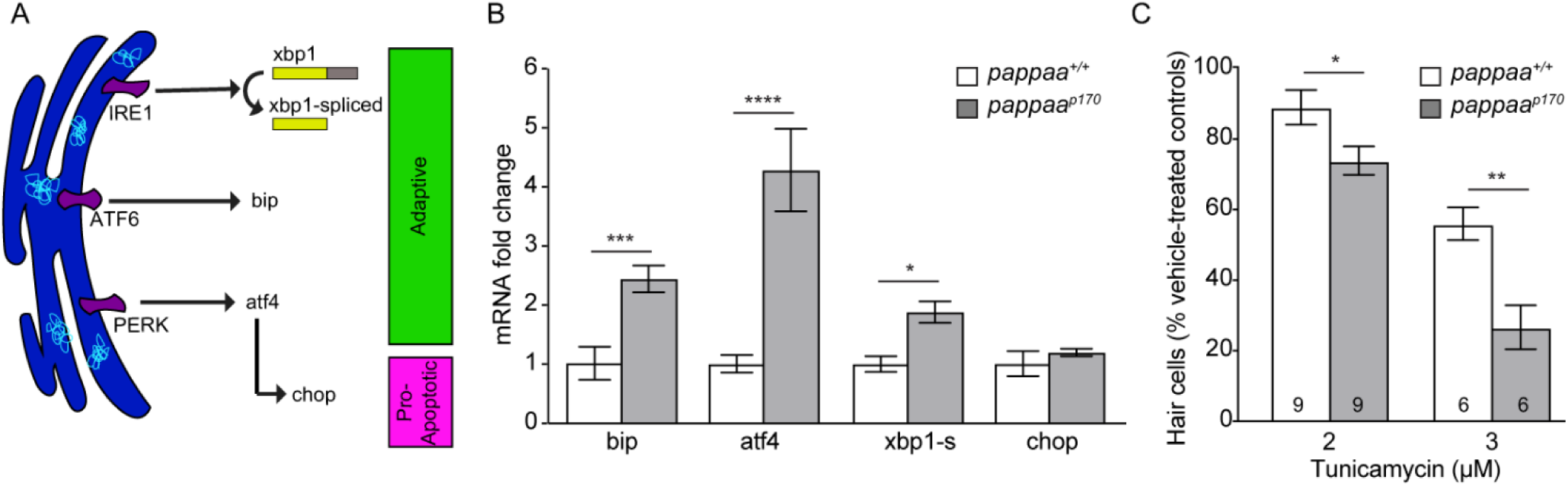
Pappaa loss causes ER stress. (A) Schematic of the UPR pathway. The accumulation of unfolded proteins activates the UPR receptors, IRE1, ATF6, and PERK, signifying ER stress. In the early adaptive phase of ER stress, the UPR promotes cell survival through the upregulation of pro-survival factors including *bip, atf4*, and *spliced xbp1*. A switch from an adaptive to a pro-apoptotic UPR occurs during the late phase of ER stress in which Chop, a pro-apoptotic transcription factor, is upregulated. (B) Mean fold change in UPR mRNA levels in wild type and *pappaa*^*p170*^ hair cells at 5 dpf. N=2–3 technical replicates/gene. \**p*<0.05, \*\*\**p*<0.001, \*\*\*\**p*<0.0001. two-way ANOVA, Holm-Sidak post test. Error bars=SEM. (C) Mean percentage of surviving hair cells following a 24-hour treatment with tunicamycin starting from 4 dpf. To calculate hair cell survival percentage, hair cell number post-treatment was normalized to mean hair cell number in vehicle treated larvae of the same genotype. \**p*<0.05, \*\**p*<0.01, two-way ANOVA, Holm-Sidak post test. N=6-9 larvae per group (shown at base of bars), 3 neuromasts/ larva from 2 experiments were analyzed. Error bars=SEM. **Figure 5-source data 1**. Mean fold change in UPR transcript levels in wild type and *pappaa*^*p170*^ lateral line hair cells. **Figure 5-source data 2**. Hair cell survival following treatment with Tunicamycin in wild type and *pappaa*^*p170*^ larvae.

## Conclusion

The ER and mitochondria are physically and functionally linked together to facilitate a range of cellular processes essential for neuron survival. Disruptions to this connection can yield damage to vital cellular components and is found to be the common denominator in neurodegenerative disease. To identify potential targets for therapeutic interventions, we must define and characterize the genetic and molecular mechanisms regulating the ER-mitochondria axis. Here, we describe a novel role for Pappaa, a local stimulator of IGF1 signaling, in regulating the ER-mitochondria axis and its essential downstream processes. We provide evidence that loss of Pappaa, and presumably the reduction of local IGF1 bioavailability, causes ER stress and excessive and abnormally tight ER-mitochondria associations resulting in mitochondrial dysfunction, reduced autophagic activity, and oxidative stress. IGF1 emerges as a promising therapeutic candidate due to its known role in influencing the function of the ER and the mitochondria. Studies using exogenous IGF1 treatment showed it can alleviate pharmacologically-induced ER stress in cancer cells (Novosyadlyy et al., 2008), or attenuate mitochondria-generated ROS in cirrhosis or aging disease models (Castilla-Cortazar et al., 1997; García-Fernández et al., 2008; Novosyadlyy et al., 2008; Sádaba et al., 2016), supporting the possibility that IGF1 signaling may play a role in regulating the ER-mitochondria axis. However, exogenous IGF1 treatment presents its own set of challenges for developing an IGF-based therapy. IGF1 signaling has a broad cellular impact and ubiquitous overexpression could affect many systems. Additionally, the suppressive effects of IGFBPs on IGF1 may render this approach inefficient. Indeed, patients with neurodegenerative disorders have not shown significant improvement following systemic IGF1 administration. These disappointing outcomes are thought to be due to the suppressive effects of IGFBPs on IGF1 bioavailability (Raoul and Aebischer, 2004; Sakowski et al., 2009). As an upstream regulator of IGF1 signaling that is spatially restricted, Pappaa may provide a more viable therapeutic alternative.

## Materials and Methods

### Maintenance of zebrafish

To generate *pappaa*^*+/+*^ and *pappaa*^*p170*^ larvae, adult *pappaa*^*p170/+*^ zebrafish (on a mixed Tubingen long-fin (*TLF*), *WIK* background) were crossed into *Tg(brn3c:mGFP)*^*s356t*^ and then incrossed. Embryonic and larval zebrafish were raised in E3 media (5 mM NaCl, 0.17 mM KCl, 0.33 mM CaCl2, 0.33 mM MgSO4, pH adjusted to 6.8–6.9 with NaHCO3) at 29°C on a 14 hr/10 hr light/dark cycle through 5 days post fertilization (dpf) (Kimmel et al., 1995; Gyda et al., 2012). All experiments were done on larvae at 5 dpf. Genotyping of *pappaa*^*p170*^ larvae was performed as previously described (Wolman et al., 2015).

### Pharmacology

All pharmacological experiments were performed on *Tg(brn3c:mGFP)* larave. Compounds were added to the larval E3 media at 4-5 dpf. Adenophostin A (Fisher Scientific 11-550-050UG; dissolved in DMSO) and Thapsigargin (Tocris No. 1138; dissolved in DMSO) were added at 10 µM for 1 hr at 5 dpf. Tunicamycin (Caymen Chemical 11445; dissolved in DMSO) was added at 2–3 µM for 24 hr, beginning at 4 dpf. Following each treatment period, larvae were washed 3 times with E3 and left to recover in E3 for 4 hr at 29°C before fixation with 4% paraformaldehyde (diluted to 4% w/v in PBS from 16% w/v in 0.1M phosphate buffer, pH 7.4). Vehicle-treated controls were treated with 0.1% DMSO.

### Immunohistochemistry

To visualize hair cells, 5 dpf larvae were fixed in 4% paraformaldehyde for 1 hr at room temperature then rinsed 3 times with PBS. Larvae were blocked for 1 hr at room temperature in incubation buffer (IB: 0.2% bovine serum albumin, 2% normal goat serum, 0.8% Triton-X, 1% DMSO, in PBS, pH 7.4). Larvae were incubated in primary antibodies (anti-GFP, 1:500, rabbit polyclonal; ThermoFisher Scientific, RRID:AB_221569) in IB overnight at 4°C. Larvae were then rinsed 5 times for 10 minutes with IB, incubated in AlexaFluor-488-conjugated secondary antibody (goat anti-rabbit polyclonal, 1:500; ThermoFisher Scientific, RRID:AB_2576217) in IB for 4 hr at room temperature, and rinsed 5 times with IB. Larvae were mounted in 70% glycerol in PBS. Images were acquired with an Olympus Fluoview confocal laser scanning microscope (FV1000) using Fluoview software (FV10-ASW 4.2).

### Hair cell survival

Hair cell survival experiments were performed in *Tg(brn3c:mGFP)* larvae as previously described (Alassaf et al., 2019). Hair cells from 3 stereotypically positioned neuromasts (IO3, M2, and OP1) (Raible and Kruse, 2000) were counted and averaged for each larva. Hair cell survival was calculated as a percentage with the following formula: [(mean number of hair cells after treatment)/ (mean number of hair cells in vehicle treated group)] X 100.

### RT-qPCR

Fluorescent sorting of *brn3c:mGFP* hair cells, RNA extraction, cDNA synthesis, and RT-qPCR were performed as previously described (Alassaf et al., 2019). The primers sequences for the UPR genes are as follows: for bip, forward: ATCAGATCTGGCCAAAATGC and reverse: CCACGTATGACGGAGTGATG; atf4, forward: TTAGCGATTGCTCCGATAGC and reverse: GCTGCGGTTTTATTCTGCTC; for chop, forward: ATATACTGGGCTCCGACACG and reverse: GATGAGGTGTTCTCCGTGGT (Howarth et al., 2014); for xbp1-spliced, forward: TGTTGCGAGACAAGACGA and reverse: CCTGCACCTGCTGCGGACT (Vacaru et al., 2014); For b-actin (endogenous control), forward TACAGCTTCACCACCACAGC and reverse: AAGGAAGGCTGGAAGAGAGC (Wang et al., 2005). Cycling conditions were as follows: 1 min at 95°C, then 40 cycles of 15 s at 95°C, followed by 1 min at 60°C (Jin et al., 2010). Analysis of relative gene expression was done using the 2−ΔΔCt method (Livak and Schmittgen, 2001).

### MitoTracker

To assess mitochondrial morphology, 5 dpf larvae were incubated in 100 nM mitotracker green FM (Thermofischer Scientific M7514; dissolved in anhydrous DMSO) for 5 min. Following the incubation period, larvae were washed three times in E3, anesthetized in 0.002% tricaine (Sigma-Aldrich) in E3, and mounted as previously described (Stawicki et al., 2014). Z-stack images were acquired at 60X with an Olympus Fluoview confocal laser scanning microscope (FV1000) using Fluoview software (FV10-ASW 4.2). A maximum intensity projection of Z-stacks that covered the full depth of the hair cells were used for analysis. Mitochondrial circularity was measured using the publicly available “mitochondrial morphology” ImageJ plugin (Dagda et al., 2009). Neomycin-Texas Red

To examine autophagy, Conjugation of Texas Red-X-succinimidyl ester to neomycin sulfate hydrate (Sigma-Aldrich) was done following the previously described protocol (Stawicki et al., 2014). A final concentration of 10 μM of neomycin-Texas Red was used. Larvae were anesthetized in 0.002% tricaine (Sigma-Aldrich) in E3, and mounted as previously described (Stawicki et al., 2014). Z*-*stack images of each neuromast were acquired at baseline (prior to Neo-TR exposure) for 2.5 minutes at 30 second intervals. Following the addition of neomycin-Texas Red to the larval media, images were acquired at 30 second intervals for 1 hour. Hair cells within each neuromast that were visually accessible with no significant physical overlap with neighboring hair cells were selected for analysis. Neomycin-Texas Red puncta were counted manually for each hair cell. Time lapse movies were made using ImageJ and corrected for xy drift using the ImageJ plugins StackReg and TurboReg (Thévenaz et al., 1998).

### Electron microscopy

5 dpf *pappaa*^*+/+*^ and *pappaa*^*p170*^ larvae in a Tubingen long-fin (*TLF*) background were processed as previously described (Miller et al., 2018). Images were acquired with a Philips CM120 scanning transmission electron microscope with a BioSprint 12 series digital camera using AMT Image Capture Software Engine V700 (Advanced Microscopy Techniques). All images were of head neuromasts. Mitochondrial morphology and ER-mitochondria associations were analyzed using ImageJ (Schneider et al., 2012). For mitochondrial morphology, a ROI was drawn around each mitochondrion using the free hand tool. The “Analyze Particles” function was used to measure the area, perimeter, and circularity. Mitochondrial interconnectivity was calculated as the area/perimeter (Wiemerslage and Lee, 2016). To measure the inter-organelle distance, the line segment tool was used to connect the 2 closest points between the ER and mitochondria. The frequency of ER-mitochondria associations was determined by counting the number of ER fragments that were within 100 nm of each mitochondrion.

### Statistics

All data were analyzed using GraphPad Prism Software 7.0b (GraphPad Software Incorporated, La Jolla, Ca, USA, RRID:SCR_002798). Data are presented as the mean ± standard error of the mean (SEM). The statistical tests and sample size for each experiment are stated in the figure legends. Significance was set at p<0.05. All data presented are from individual experiments expect for data in Figure 5C. Data collected from multiple experiments were normalized to their respective controls.

## Contribution

Conceptualization, Investigation, Data curation, Formal analysis, Writing – original draft preparation.

## Contribution

Resources, Project administration, Writing—review and editing

## Funding

*UW-Madison graduate support funds*

- Mary Halloran

*Ministry of Education-Saudi Arabia*

- Mroj Alassaf

## Acknowledgments

The authors would like to thank Benjamin August (UW medical school electron microscope facility) for his technical expertise, Dr. Yevgenya Grinblat (University of Wisconsin-Department of Integrative Biology) for the use of the RT-qPCR cycler, Emily Daykin for her assistance with data acquisition, and Daniel North for fish care and maintenance.

## Ethics

This study was performed in accordance with the recommendations in the Guide for the Care and Use of Laboratory Animals of the National Institutes of Health. Animals were handled according to approved institutional animal care and use committee (IACUC) protocols (L005704) of the University of Wisconsin.

## References

Adam-Vizi V, Starkov AA (2010) Calcium and mitochondrial reactive oxygen species generation: how to read the facts. J Alzheimers Dis 20 Suppl 2:S413–426.

Alassaf M, Daykin EC, Mathiaparanam J, Wolman MA (2019) Pregnancy-associated plasma protein-aa supports hair cell survival by regulating mitochondrial function. Elife 8.

Area-Gomez E, Del Carmen Lara Castillo M, Tambini MD, Guardia-Laguarta C, de Groof AJ, Madra M, Ikenouchi J, Umeda M, Bird TD, Sturley SL, Schon EA (2012) Upregulated function of mitochondria-associated ER membranes in Alzheimer disease. EMBO J 31:4106–4123.

Arruda AP, Pers BM, Parlakgul G, Guney E, Inouye K, Hotamisligil GS (2014) Chronic enrichment of hepatic endoplasmic reticulum-mitochondria contact leads to mitochondrial dysfunction in obesity. Nat Med 20:1427–1435.

Bernard-Marissal N, Médard JJ, Azzedine H, Chrast R (2015) Dysfunction in endoplasmic reticulum-mitochondria crosstalk underlies SIGMAR1 loss of function mediated motor neuron degeneration. Brain 138:875–890.

Bravo R, Gutierrez T, Paredes F, Gatica D, Rodriguez AE, Pedrozo Z, Chiong M, Parra V, Quest AF, Rothermel BA, Lavandero S (2012) Endoplasmic reticulum: ER stress regulates mitochondrial bioenergetics. Int J Biochem Cell Biol 44:16–20.

Bravo R, Vicencio JM, Parra V, Troncoso R, Munoz JP, Bui M, Quiroga C, Rodriguez AE, Verdejo HE, Ferreira J, Iglewski M, Chiong M, Simmen T, Zorzano A, Hill JA, Rothermel BA, Szabadkai G, Lavandero S (2011) Increased ER-mitochondrial coupling promotes mitochondrial respiration and bioenergetics during early phases of ER stress. J Cell Sci 124:2143–2152.

Brookes PS, Yoon Y, Robotham JL, Anders MW, Sheu SS (2004) Calcium, ATP, and ROS: a mitochondrial love-hate triangle. Am J Physiol Cell Physiol 287:C817–833.

Calì T, Ottolini D, Negro A, Brini M (2013) Enhanced parkin levels favor ER-mitochondria crosstalk and guarantee Ca(2+) transfer to sustain cell bioenergetics. Biochim Biophys Acta 1832:495–508.

Castilla-Cortazar I, Garcia M, Muguerza B, Quiroga J, Perez R, Santidrian S, Prieto J (1997) Hepatoprotective effects of insulin-like growth factor I in rats with carbon tetrachloride-induced cirrhosis. Gastroenterology 113:1682–1691.

Chaudhari N, Talwar P, Parimisetty A, Lefebvre d’Hellencourt C, Ravanan P (2014) A molecular web: endoplasmic reticulum stress, inflammation, and oxidative stress. Front Cell Neurosci 8:213.

Csordás G, Weaver D, Hajnóczky G (2018) Endoplasmic Reticulum-Mitochondrial Contactology: Structure and Signaling Functions. Trends Cell Biol 28:523–540.

Csordás G, Renken C, Várnai P, Walter L, Weaver D, Buttle KF, Balla T, Mannella CA, Hajnóczky G (2006) Structural and functional features and significance of the physical linkage between ER and mitochondria. J Cell Biol 174:915–921.

Cárdenas C, Miller RA, Smith I, Bui T, Molgó J, Müller M, Vais H, Cheung KH, Yang J, Parker I, Thompson CB, Birnbaum MJ, Hallows KR, Foskett JK (2010) Essential regulation of cell bioenergetics by constitutive InsP3 receptor Ca2+ transfer to mitochondria. Cell 142:270–283.

Deniaud A, Sharaf el dein O, Maillier E, Poncet D, Kroemer G, Lemaire C, Brenner C (2008) Endoplasmic reticulum stress induces calcium-dependent permeability transition, mitochondrial outer membrane permeabilization and apoptosis. Oncogene 27:285–299.

Ding Y, Li J, Liu Z, Liu H, Li H, Li Z (2017) IGF-1 potentiates sensory innervation signalling by modulating the mitochondrial fission/fusion balance. Sci Rep 7:43949.

Eggermont JJ (2017) Hearing loss : causes, prevention, and treatment. In, pp 1 online resource (xxi, 391 pages). London, U.K.: Academic Press is an imprint of Elsevier,.

Esterberg R, Hailey DW, Rubel EW, Raible DW (2014) ER-mitochondrial calcium flow underlies vulnerability of mechanosensory hair cells to damage. J Neurosci 34:9703–9719.

Friedman JR, Lackner LL, West M, DiBenedetto JR, Nunnari J, Voeltz GK (2011) ER tubules mark sites of mitochondrial division. Science 334:358–362.

Fu S, Yalcin A, Lee GY, Li P, Fan J, Arruda AP, Pers BM, Yilmaz M, Eguchi K, Hotamisligil GS (2015) Phenotypic assays identify azoramide as a small-molecule modulator of the unfolded protein response with antidiabetic activity. Sci Transl Med 7:292ra298.

García-Fernández M, Delgado G, Puche JE, González-Barón S, Castilla Cortázar I (2008) Low doses of insulin-like growth factor I improve insulin resistance, lipid metabolism, and oxidative damage in aging rats. Endocrinology 149:2433–2442.

Gomez-Suaga P, Paillusson S, Miller CCJ (2017a) ER-mitochondria signaling regulates autophagy. Autophagy 13:1250–1251.

Gomez-Suaga P, Paillusson S, Stoica R, Noble W, Hanger DP, Miller CCJ (2017b) The ER-Mitochondria Tethering Complex VAPB-PTPIP51 Regulates Autophagy. Curr Biol 27:371–385.

Gonzalez-Gonzalez S (2017) The role of mitochondrial oxidative stress in hearing loss. Neurological Disorders and Therapeutics 1:1–5.

Gorlach A, Bertram K, Hudecova S, Krizanova O (2015) Calcium and ROS: A mutual interplay. Redox Biol 6:260–271.

Gregersen N, Bross P, Vang S, Christensen JH (2006) Protein misfolding and human disease. Annu Rev Genomics Hum Genet 7:103–124.

Hailey DW, Esterberg R, Linbo TH, Rubel EW, Raible DW (2017) Fluorescent aminoglycosides reveal intracellular trafficking routes in mechanosensory hair cells. J Clin Invest 127:472–486.

He Z, Guo L, Shu Y, Fang Q, Zhou H, Liu Y, Liu D, Lu L, Zhang X, Ding X, Tang M, Kong W, Sha S, Li H, Gao X, Chai R (2017) Autophagy protects auditory hair cells against neomycin-induced damage. Autophagy 13:1884–1904.

Hedskog L, Pinho CM, Filadi R, Rönnbäck A, Hertwig L, Wiehager B, Larssen P, Gellhaar S, Sandebring A, Westerlund M, Graff C, Winblad B, Galter D, Behbahani H, Pizzo P, Glaser E, Ankarcrona M (2013) Modulation of the endoplasmic reticulum-mitochondria interface in Alzheimer’s disease and related models. Proc Natl Acad Sci U S A 110:7916–7921.

Houck SA, Singh S, Cyr DM (2012) Cellular responses to misfolded proteins and protein aggregates. Methods Mol Biol 832:455–461.

Hwa V, Oh Y, Rosenfeld RG (1999) The insulin-like growth factor-binding protein (IGFBP) superfamily. Endocr Rev 20:761–787.

Iqbal S, Hood DA (2014) Oxidative stress-induced mitochondrial fragmentation and movement in skeletal muscle myoblasts. Am J Physiol Cell Physiol 306:C1176–1183.

Ježek J, Cooper KF, Strich R (2018) Reactive Oxygen Species and Mitochondrial Dynamics: The Yin and Yang of Mitochondrial Dysfunction and Cancer Progression. Antioxidants (Basel) 7.

Krols M, Bultynck G, Janssens S (2016) ER-Mitochondria contact sites: A new regulator of cellular calcium flux comes into play. J Cell Biol 214:367–370.

Lee KS, Huh S, Lee S, Wu Z, Kim AK, Kang HY, Lu B (2018) Altered ER-mitochondria contact impacts mitochondria calcium homeostasis and contributes to neurodegeneration in vivo in disease models. Proc Natl Acad Sci U S A 115:E8844–E8853.

Liesa M, Palacín M, Zorzano A (2009) Mitochondrial dynamics in mammalian health and disease. Physiol Rev 89:799–845.

Liu Y, Zhu X (2017) Endoplasmic reticulum-mitochondria tethering in neurodegenerative diseases. Transl Neurodegener 6:21.

Marchi S, Rimessi A, Giorgi C, Baldini C, Ferroni L, Rizzuto R, Pinton P (2008) Akt kinase reducing endoplasmic reticulum Ca2+ release protects cells from Ca2+-dependent apoptotic stimuli. Biochem Biophys Res Commun 375:501–505.

McPherson DR (2018) Sensory Hair Cells: An Introduction to Structure and Physiology. Integr Comp Biol 58:282–300.

Ni HM, Williams JA, Ding WX (2015) Mitochondrial dynamics and mitochondrial quality control. Redox Biol 4:6–13.

Novosyadlyy R, Kurshan N, Lann D, Vijayakumar A, Yakar S, LeRoith D (2008) Insulin-like growth factor-I protects cells from ER stress-induced apoptosis via enhancement of the adaptive capacity of endoplasmic reticulum. Cell Death Differ 15:1304–1317.

Oslowski CM, Urano F (2011) Measuring ER stress and the unfolded protein response using mammalian tissue culture system. Methods Enzymol 490:71–92.

Rizzuto R, De Stefani D, Raffaello A, Mammucari C (2012) Mitochondria as sensors and regulators of calcium signalling. Nat Rev Mol Cell Biol 13:566–578.

Rizzuto R, Pinton P, Carrington W, Fay FS, Fogarty KE, Lifshitz LM, Tuft RA, Pozzan T (1998) Close contacts with the endoplasmic reticulum as determinants of mitochondrial Ca2+ responses. Science 280:1763–1766.

Rowland AA, Voeltz GK (2012) Endoplasmic reticulum-mitochondria contacts: function of the junction. Nat Rev Mol Cell Biol 13:607–625.

Sano R, Hou YC, Hedvat M, Correa RG, Shu CW, Krajewska M, Diaz PW, Tamble CM, Quarato G, Gottlieb RA, Yamaguchi M, Nizet V, Dahl R, Thomas DD, Tait SW, Green DR, Fisher PB, Matsuzawa S, Reed JC (2012) Endoplasmic reticulum protein BI-1 regulates Ca^2+^-mediated bioenergetics to promote autophagy. Genes Dev 26:1041–1054.

Sádaba MC, Martín-Estal I, Puche JE, Castilla-Cortázar I (2016) Insulin-like growth factor 1 (IGF-1) therapy: Mitochondrial dysfunction and diseases. Biochim Biophys Acta 1862:1267–1278.

Urra H, Hetz C (2012) The ER in 4D: a novel stress pathway controlling endoplasmic reticulum membrane remodeling. Cell Death Differ 19:1893–1895.

Wiemerslage L, Lee D (2016) Quantification of mitochondrial morphology in neurites of dopaminergic neurons using multiple parameters. J Neurosci Methods 262:56–65.

Zhu G, Lee AS (2015) Role of the unfolded protein response, GRP78 and GRP94 in organ homeostasis. J Cell Physiol 230:1413–1420.

